# Identifying miRNA-mRNA regulatory relationships in breast cancer with invariant causal prediction

**DOI:** 10.1101/340638

**Authors:** Vu Viet Hoang Pham, Junpeng Zhang, Lin Liu, Buu Minh Thanh Truong, Taosheng Xu, Trung Tin Nguyen, Jiuyong Li, Thuc Duy Le

## Abstract

microRNAs (miRNAs) regulate gene expression at the post-transcriptional level and they play an important role in various biological processes in the human body. Therefore, identifying their regulation mechanisms is essential for the diagnostics and therapeutics for a wide range of diseases. There have been a large number of researches which use gene expression profiles to resolve this problem. However, the current methods have their own limitations. Some of them only identify the correlation of miRNA and mRNA expression levels instead of the causal or regulatory relationships while others infer the causality but with a high computational complexity. To overcome these issues, in this study, we propose a method to identify miRNA-mRNA regulatory relationships in breast cancer using the invariant causal prediction. The key idea of invariant causal prediction is that the cause miRNAs of their target mRNAs are the ones which have persistent causal relationships with the target mRNAs across different environments. In this research, we aim to find miRNA targets which are consistent across different breast cancer subtypes. Thus, first of all, we apply the Pam50 method to categorise BRCA samples into different ‘‘environment” groups based on different cancer subtypes. Then we use the invariant causal prediction method to find miRNA-mRNA regulatory relationships across subtypes. We validate the results with the miRNA-transfected experimental data and the results show that our method outperforms the state-of-the-art methods. In addition, we also integrate this new method with the Pearson correlation analysis method and Lasso in an ensemble method to take the advantages of these methods. We then validate the results of the ensemble method with the experimentally confirmed data and the ensemble method shows the best performance, even comparing to the proposed causal method. Functional enrichment analyses show that miRNAs in the regulatory relationship predicated by the proposed causal method tend to synergistically regulate target genes, indicating the usefulness of these methods, and the identified miRNA targets could be used in the design of wet-lab experiments to discover the causes of breast cancer.

**Author summary:** Cancer is a disease of cells in human body and it causes a high rate of deaths world wide. There has been evidence that non-coding RNAs are key players in the development and progression of cancer. Among the different types of non-coding RNAs, miRNAs, which are short non-coding RNAs, regulate gene expression and play an important role in different biological processes as well as various cancer types. To design better diagnostic and therapeutic plans for cancer patients, we need to know the roles of miRNAs in cancer initialisation and development, and their regulation mechanisms in the human body. In this study, we propose algorithms to identify miRNA-mRNA regulatory relationships in breast cancer. Comparing our methods with existing methods in predicting miRNA targets, our methods show a better performance. The estimated miRNA targets from our methods could be a potential source for further wet-lab experiments to discover the causes of breast cancer.

## Introduction

The human transcriptome is composed of 98% of non-coding RNAs (ncRNAs) and only 2% of protein-coding RNAs [1]. However, research into the roles of ncRNAs is still in the early stage. The emergence of ncRNAs as new key players in cancer development and progression has shifted our understanding of gene regulation [1, 2], especially since the discovery of microRNAs (miRNAs). miRNAs are short ncRNAs that regulate gene expression at the post-transcriptional level and identified as the drivers in diverse disease conditions including cancers, where they function either as oncogenes or as tumour suppressors [3, 4]. Recent years have also seen the discovery of several other types of ncRNAs, including long non-coding RNAs (lnRNAs), pseudogenes and circular RNAs (cirRNAs), along with their regulatory functions in disease conditions [4]. There also has been evidence that mRNAs, miRNAs, and other ncRNAs work in concert to regulate cancer development and progression [5, 6].

There have been several methods developed to explore miRNA functions, including those for predicting miRNA targets and regulatory modules (see [7] for a review), inferring miRNA sponge networks and modules [6,8–10], and identifying cancer subtypes [11–13]. However, our understanding of miRNAs’ roles in regulating cancer across different subtypes thereby permitting prognosis, diagnosis, and prediction of therapy response is still very far from complete, and reliable methods for identifying miRNA-mRNA regulatory relationships in cancer are in demand.

Existing computational methods for inferring miRNA-mRNA regulatory relationships are of two major categories: sequence based approach and expression based approach. The former is based on complementary base pairing, site accessibility, and evolutionary conservation; and the latter relies on the negative correlation between miRNA and mRNA expression levels. This expression-based approach can be further divided into i) correlation-based approach [14–16], and ii) causal inference approach [17–19].

Each of the approaches has its own advantages and limitations. The correlation and regression based approaches [14–16] are efficient for large gene expression datasets. However, correlations or associations are not causality, but miRNA-mRNA regulatory relationships are causal relationships. A strong correlation between the expression values of a miRNA and a mRNA in a dataset may be a spurious relationship, as it could be confounded by a transcription factor. On the other hand, causal inference approach [17–19] aims to estimate the intervention effects as in the gene knockdown experiments. While these causal inference methods help remove spurious relationships, they have high computational complexity and therefore they are not scalable in large datasets. Moreover, these methods do causal inference based on the causal graphs learnt from data, which involves false discoveries when the sample size is not large enough.

In this paper, we propose to infer the miRNA-mRNA regulatory relationships in breast cancer by adapting a recently developed causal inference method, invariant causal prediction (ICP) [20]. Applying the key idea of causal invariance used by ICP, the causes (miRNAs) of a mRNA are the ones that show consistent causal relationships with the mRNA across different environments. The “different environments” can be understood as different datasets obtained from different sources/labs for studying the same disease, or different types of datasets such as observational data and data obtained from intervention experiments. In this paper, we aim to find the miRNA-mRNA regulatory relationships that are persistent across different subtypes of breast cancer. We firstly apply the PAM50 method [21] to the TCGA breast cancer dataset to classify the samples into 5 different breast cancer subtypes (Basal, Her2, LumA, LumB, Normal-like). We then use ICP to search for miRNA-mRNA pairs that show persistent causal relationship across different subtypes. Validating the predictions with miRNA transfection data, the results show that our proposed method performs better than the exising methods that are based on correlation, regression and other causal discovery methods such as idaFast or jointIDA. Furthermore, the method is much faster than the other existing causal discovery based methods. We also develop an ensemble method that combines the proposed method with a correlation based method (Pearson) and a regression based method (Lasso) to take the merits of different approaches. Using experimentally confirmed databases, miRTarbase 6.1, TarBase 7.0 and miRWalk 2.0, we show that the ensemble method is the best method compared to its individual component methods, including the proposed causal invariance method. In addition, functional enrichment analyses show that the identified miRNA-mRNA relationships are highly enriched in functions and processes related to breast cancer, suggesting the usefulness of the method. Novel interactions identified by the proposed methods could be good candidates for follow-up wet-lab experiments to explore their roles in breast cancer.

## Methods

### Overview

The overview of our method is in Fig 1. It has three main steps, including selecting significant miRNAs and mRNAs, categorising samples into different experiment settings and predicting causal effects of miRNAs on mRNAs. The detail of the method is described in the following sections.

**Fig 1.**
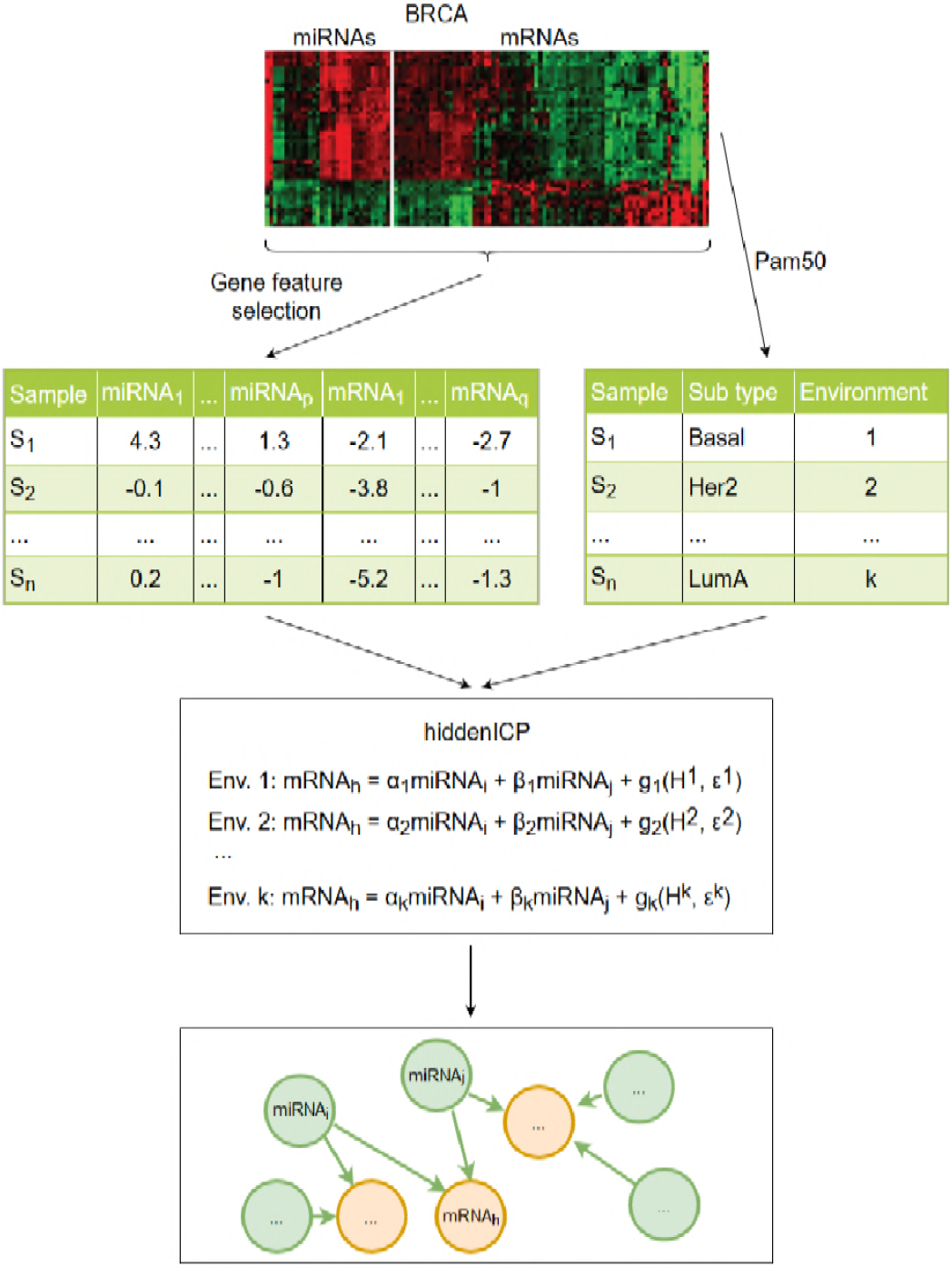
The overview of our method. The method includes three main steps, i) Select significant miRNAs and mRNAs, ii) Categorise samples into different experiment settings and iii) Predict causal effects of miRNAs on mRNAs.

### Procedure of identifying miRNA-mRNA regulatory relationships in cancer using hidden invariant causal prediction

The algorithm for detecting miRNA-mRNA relationships includes three steps as the followings.

**Step 1**: Select significant miRNAs and mRNAs. The matched miRNA and mRNA expression samples are extracted from the breast adenocarcinoma (BRCA) dataset of The Cancer Genome Atlas (TCGA) [22]. In total 503 samples with matched miRNA and mRNA expression are obtained and stored in S1 File. Then we use the FSbyMAD function of CancerSubtypes package [11] to select miRNAs and mRNAs with the most different Median Absolute Deviation (MAD). We select the top 30 miRNAs and top 1,500 mRNAs for our experiments so that other causal inference methods including jointIDA or IDA could produce the results within a week for the purpose of comparison.

**Step 2**: Categorise samples into different experiment settings based on cancer subtypes by using Pam50 [21] to discover miRNA targets across cancer subtypes.

**Step 3**: Estimate the causal relationship of miRNAs on mRNAs by estimating the causal relationship of miRNAs on each mRNA through hiddenICP function of InvariantCausalPrediction package [20]. The detail of this step is as the following.

#### Invariant causal predictation

The ICP method considers that the causal relationship between the target and each of its direct causes maintains invariant across different environments. Based on this causal invariance idea, ICP aims to find the complete set of parents (direct causes) of the target variable by searching for the subset of predictors such that in different environments, given this subset of predictors, the conditional probabilities of the target are the same. Below are the details of the method.

We use the similar notation as that in [20]. Let 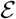 be the set of environments. For an environment 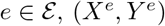 is an independent and identically distributed (i.i.d.) sample in e where *X^e^* is the set of predictor variables and *Y^e^* is the target variable. *X^e^* has *p* elements and *X^e^* ∈ ℝ^*p*^, and *Y^e^* ∈ ℝ. Let 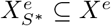 be the subset of causal predictor variables or direct causes of *Y^e^*, where 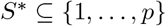 is the indices of the predictor variables, then ICP assumes the following condition holds 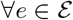:

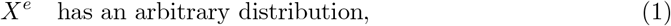

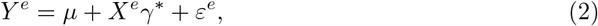

where *μ* is a constant intercept term, 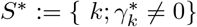, i.e. the non-zero coefficients indicate the support of the predictor variables in 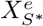, and 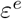 is the error distribution

The above assumption can be interpreted as 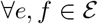 : the conditionals 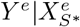, and 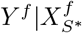, are the same.

#### Hidden invariant causal predictation

ICP has an extension for hidden variables. The hidden ICP assumes that 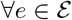:

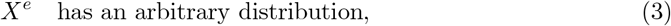

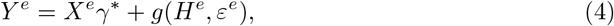

where *H* are hidden variables, *γ*\* ∈ ℝ^*p*^ are causal coefficients and *g* : ℝ^*q*^ × ℝ → ℝ is a function

In this work, we propose to apply hidden ICP to discover miRNA-mRNA regulatory relationships. This choice (instead of normal ICP) is based on the fact that in the data preparation step, we only select significant miRNAs and mRNAs as the input of ICP. Therefore in the corresponding dataset, there might be hidden miRNAs which are regulators of mRNAs. In our application of hidden ICP, the set of significant miRNAs are considered as the predictor variables. Then for each mRNA (the target or response variable), hidden ICP is used to find the direct causes, i.e. the miRNAs which regulate of the mRNA. In addition, we use Pam50 [21] to categorise the samples into different subtypes, and consider the subtypes as the environments used in hidden ICP.

### Implementation

The above algorithm has been implemented and integrated into the R package miRLAB [23]. In addition, the R scripts for reproducing the results of experiments in this paper are also available upon request.

### Functional annotation of miRNAs

We do enrichment analysis for miRNA targets to annotate biological functions of miRNAs. We apply GO [24], KEGG [25], Reactome [26] and DisGeNET [27] for the top target genes based on the point estimator for the causal effects of each miRNA identified by hiddenICP using Pam50 (hiddenICP-Pam50). Since the enrichment analysis results of hundreds of target genes are too general to gain biological insight, we only focus on the enrichment analysis of the top 50 target genes for each miRNA.

## Results and discussion

The predicted results of miRNA-mRNA regulation relationships are validated with the transfection data [23], which can be found in S4 File. In addition, the experimentally confirmed databases are also used for the validation of the results, including miRTarbase version 6.1 [28], TarBase version 7.0 [29], and miRWalk version 2.0 [30]. The confirmed miRNA-mRNA interactions retrieved from these databases are available in S5 File.

### Comparation of results

To evaluate the performance of hidden ICP, we have used the other 4 methods in our experiments for comparison, including idaFast [17] in pcalg package [31], jointIDA_direct [32], Pearson [33] and Lasso [34]. idaFast is a function which is used to estimate total causal effect of one variable on various target variables. jointIDA estimates total joint effect of a set of variables on another variable. Pearson and Lasso estimate the correlation coefficient and the regression coefficient of two variables respectively. These methods are chosen because idaFast and jointIDA are causal methods with similar goal as ours while Pearson and Lasso are popular correlation and regression methods.

With hidden IPC, we run it in two separate scenarios. In the first scenario, we randomly divide the samples into three datasets, each corresponding to an environment. In the second scenario, Pam50 [21] is used to categorise the samples based on different cancer subtypes, including Basal, Her2, LumA, LumB, and Normal-like, to create datasets for the different environments.

The top miRNA-mRNA interactions predicated by each of the 6 methods are selected to be validated with the transfection data and experimentally confirmed interactions. To have a comprehensive analysis, we select the top 500, 1,000, 1,500, and 2,0 miRNA-mRNA interactions for the validation, and we also do the validation with respect to each miRNA by selecting the top 50, 100, 150 and 200 interactions in which the miRNA is involved.

First of all, we validate the results of the 6 methods by using the transfection data as the ground truth. As the miRNAs in the transfection data are not complete, for this case, it is not fair to compare the top miRNA-mRNA interactions for all miRNAs. Thus, for the validation using the transfection data, we only compare the results based on the top of miRNA-mRNA interactions with respect to each of the miRNAs. The comparison result is shown in Fig 2. It can be seen that in all four cases with the top 50, 100, 150 and 200 ‘‘interactions predicted” for each miRNA, hiddenICP using Pam50 (hiddenICP-Pam50 in the figure) outperforms the other methods in discovering miRNA-mRNA regulation relationships. When combining with Pam50, hiddenICP (i.e. hiddenICP-Pam50) shows the best performance, indicating that the method may serve as a good tool in predicting miRNA targets. To further validate this conclusion, in the following, we analyse the results of the validation with the experimentally confirmed databases.

**Fig 2.**
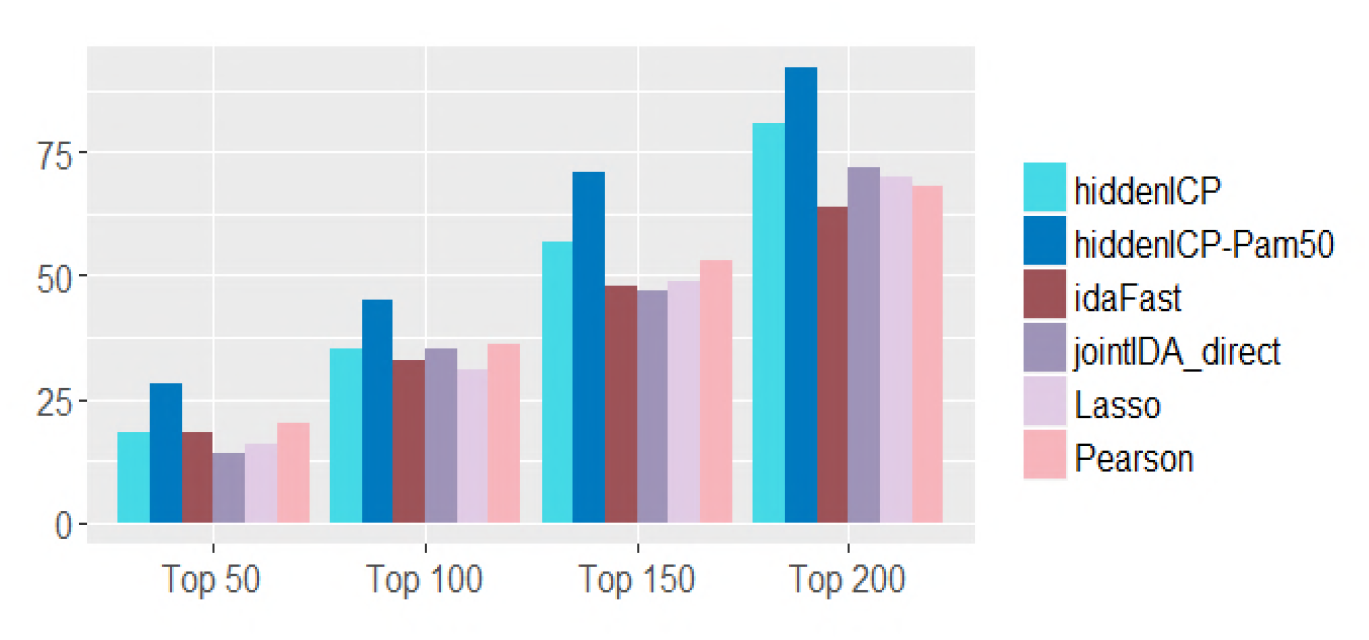
Validation using the transfection data. For each miRNA, the top 50, 100, 150 and 200 predicted miRNA-mRNA interactions are selected and validated against the transfection data. Each bar in the diagram shows the total number of validated interactions accumulated over all the miRNAs validated.

When we validate the top predicted miRNA-mRNA interactions with the experimentally confirmed databases, none of methods find a significant number of experimentally confirmed miRNA-mRNA interactions. In addition, each method uncovers some similar miRNA-mRNA pairs and some different pairs from others as in Fig 3 and Fig 4. Fig 3 shows the intersection of predicted results of methods with top 2,0 miRNA targets for all miRNAs while Fig 4 shows the intersection of predicted results of methods with top 200 miRNA targets for each miRNA. Although it looks like that Pearson and Lasso detect more interactions than other methods, we cannot have the conclusion that they are better than others as others could discover some interactions which cannot be identified by Pearson or Lasso.

**Fig 3.**
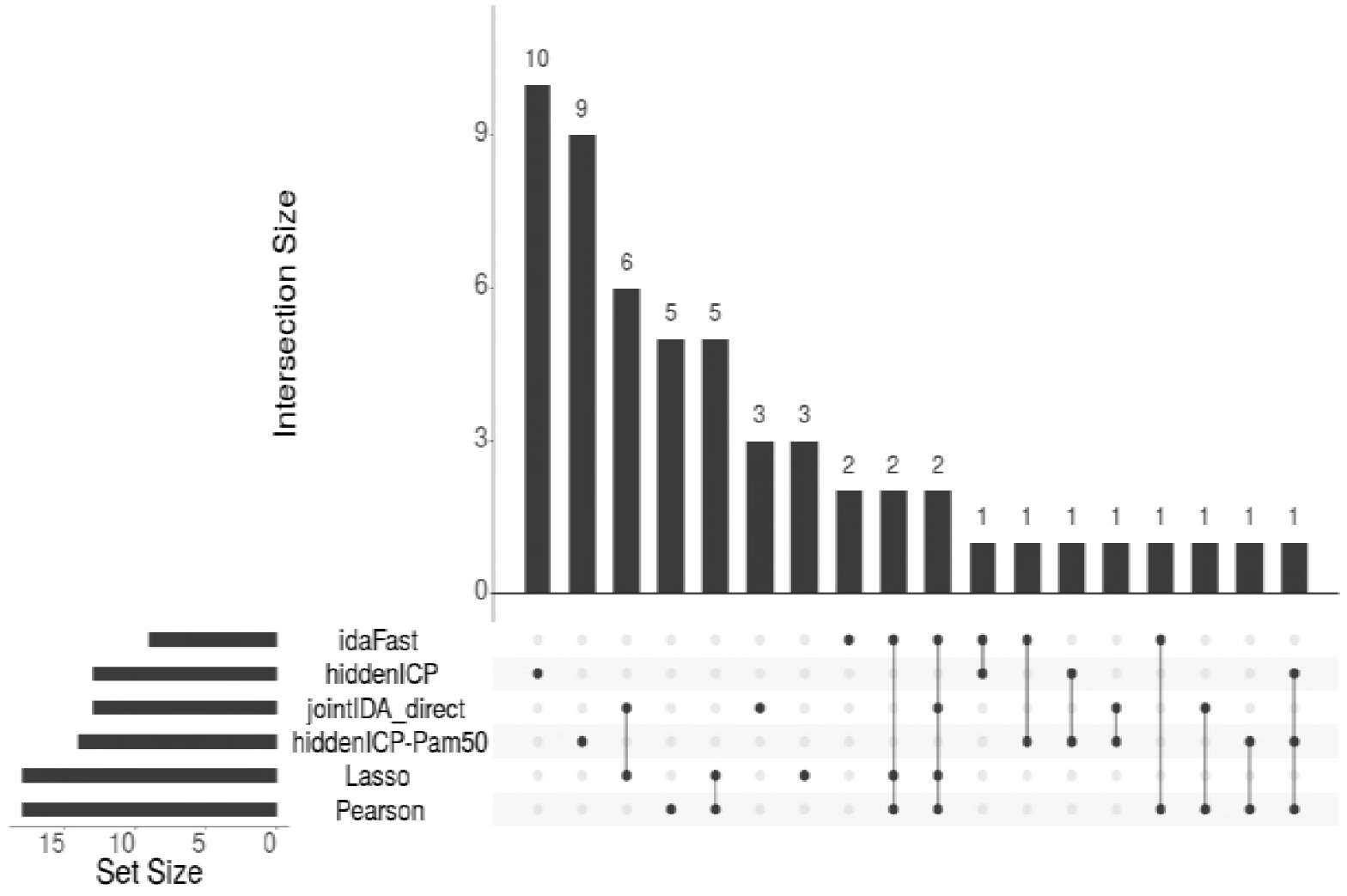
Overlap between different methods. The top miRNA-mRNA interactions validated by using the experimentally confirmed database information. For each method, the figure shows that among the top 2,000 predicted miRNA-mRNA interactions, how many interactions have been validated to be true by the databases (on the bottom left), and between the different methods how the validated interactions overlap with each other (the dotted lines and the diagram on top).

**Fig 4.**
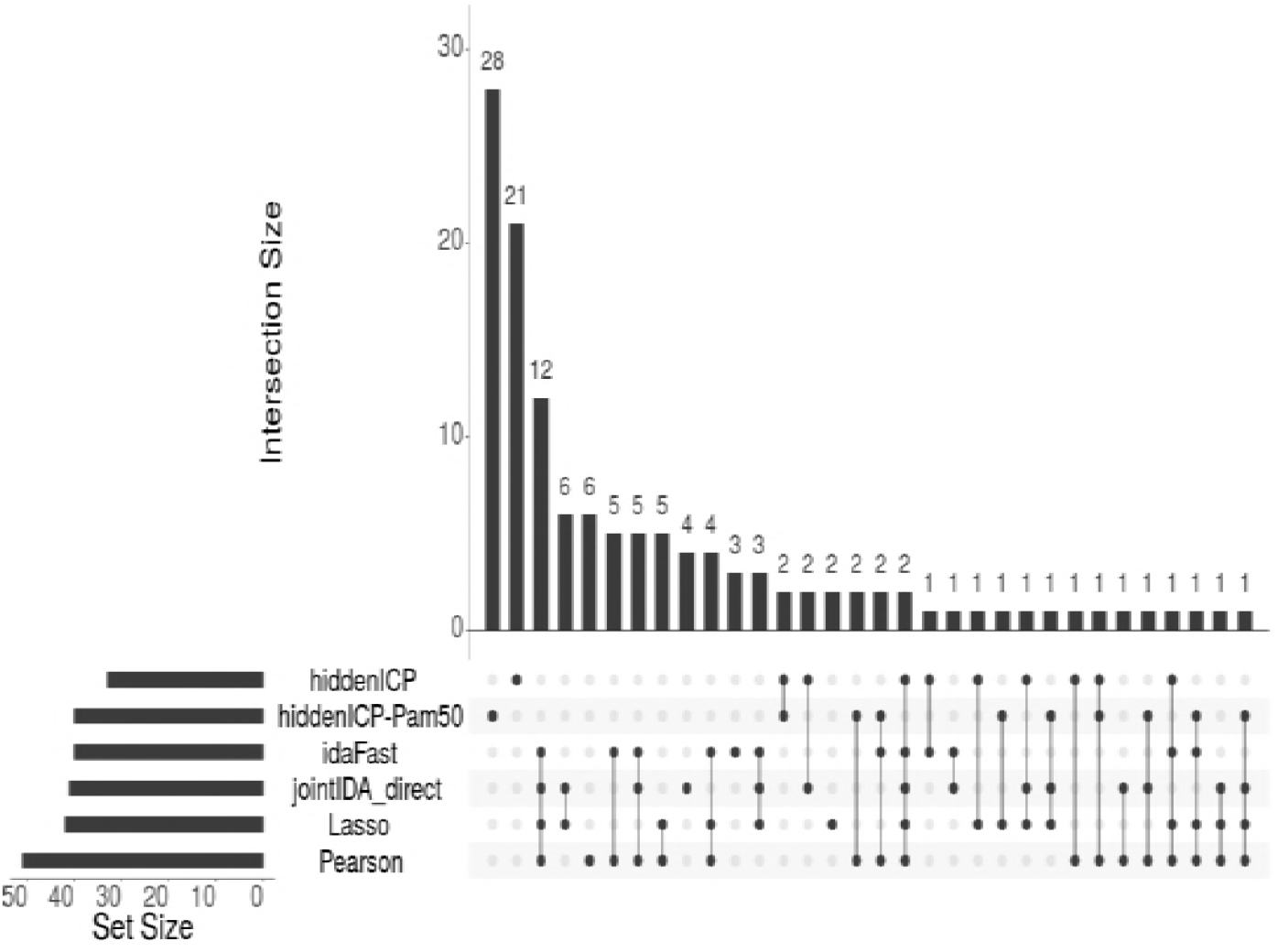
Overlap between different methods. The top miRNA-mRNA interactions validated by using the experimentally confirmed database information. For each method, the figure shows that among the top 200 predicted miRNA-mRNA interactions, how many interactions have been validated to be true by the databases (on the bottom left), and between the different methods how the validated interactions overlap with each other (the dotted lines and the diagram on top).

#### Hidden ICP forms a good performance in identifying miRNA-mRNA regulatory relationships of ensemble method

Based on the observations that different methods may provide complementary findings of miRNA-mRNA interactions, and Pearson and Lasso individually may perform better than the other methods, we use the Borda function in the package miRLAB [23] to integrate Pearson [33], Lasso [34] with others to generate ensembles for predicting miRNA-mRNA interactions. This ensemble method will get the average of the rankings from individual methods. The validation results of the ensembles are shown in Fig 5 and Fig 6, for the validation of the collection of top interactions of all miRNAs and the validation of the top interactions around individual miRNAs, respectively. In both cases, the Borda with Pearson, Lasso and hiddenICP using Pam50 outperforms others.

**Fig 5.**
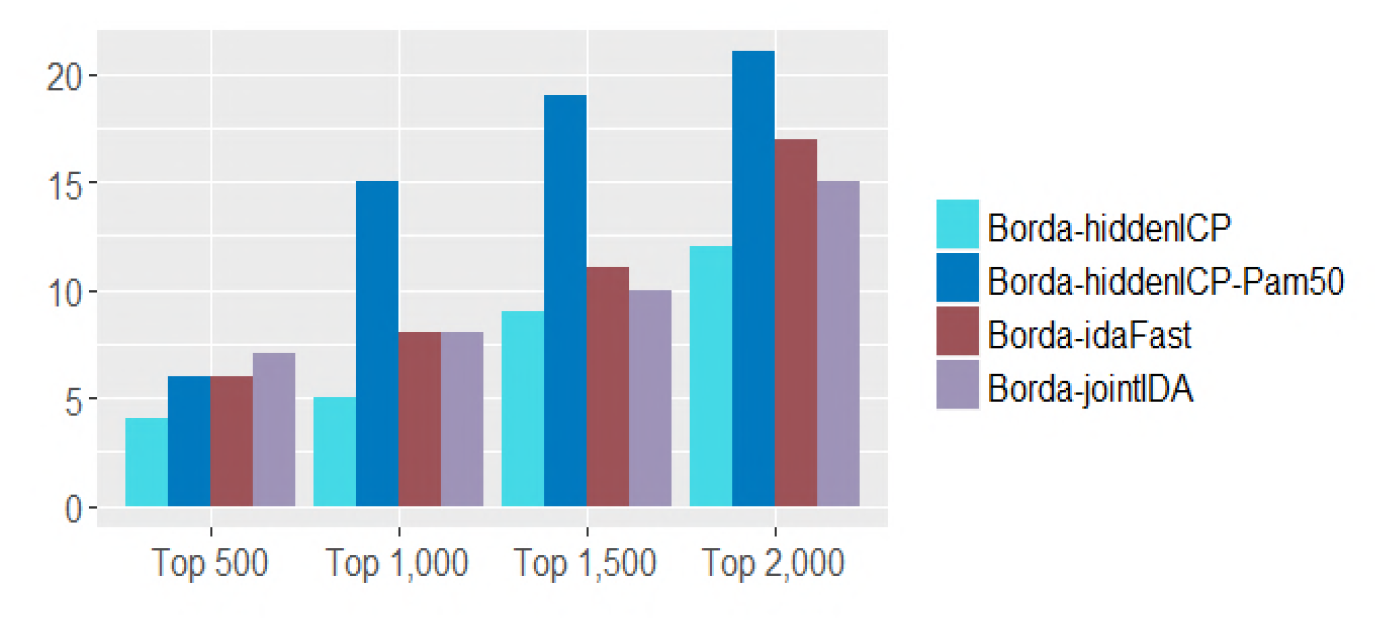
Validation using the experimentally confirmed databases. The top 500, 1,000, 1,500 and 2,000 predicted miRNA-mRNA interactions are selected and validated against the experimentally confirmed databases. Each bar in the diagram shows the total number of validated interactions.

**Fig 6.**
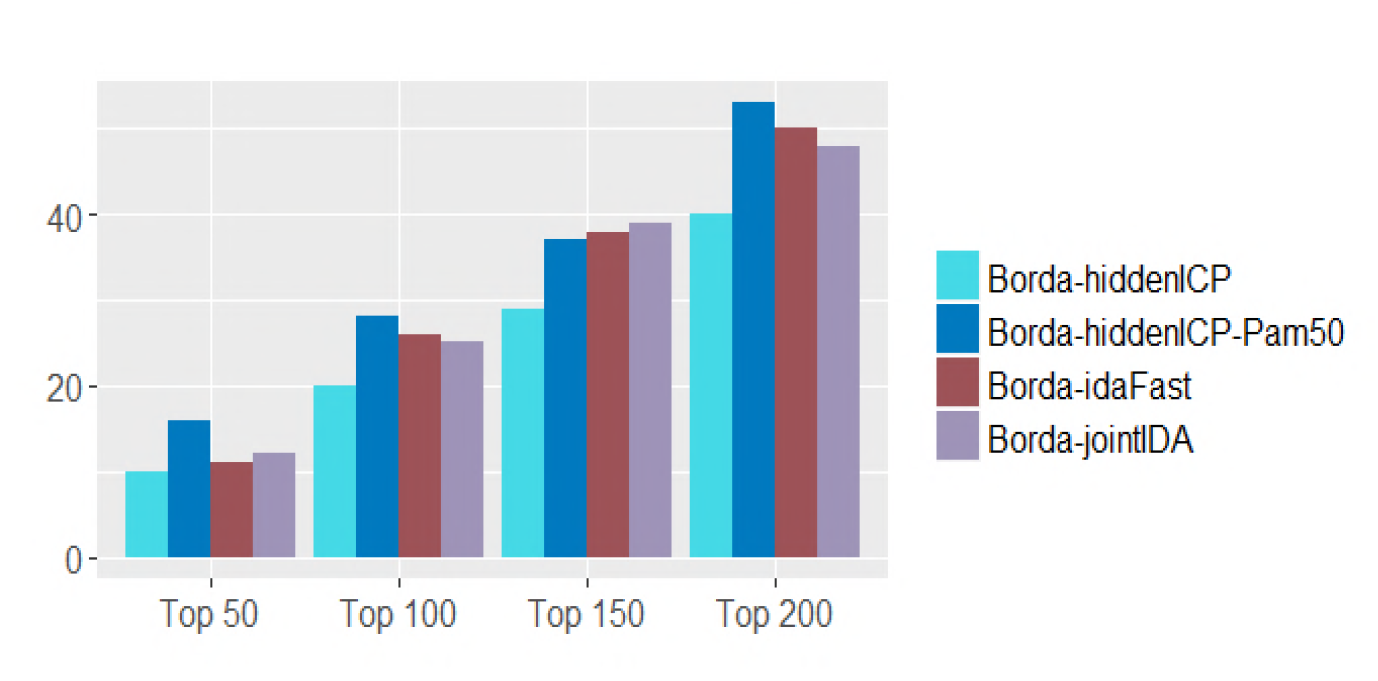
Validation using the experimentally confirmed databases. For each miRNA, the top 50, 100, 150 and 200 predicted miRNA-mRNA interactions are selected and validated against the experimentally confirmed databases. Each bar in the diagram shows the total number of validated interactions accumulated over all the miRNAs validated.

### miRNAs tend to synergistically regulate target genes

In this section, we focus on studying miRNA synergism based on the top 50, 100, 150 and 200 target genes for each miRNA identified by hiddenICP-Pam50. For each possible *miRNA* synergistic pair *miRNAi* and *miRNA_j_, i ≠ j*, the hypergeometric test is used to evaluate the significance of the shared mRNAs by these two miRNAs. The significance p-value is calculated as follows:

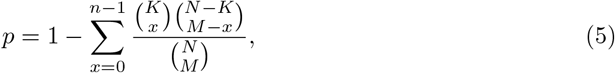

where *N* denotes the number of all mRNAs of interest, *K* is the number of mRNAs interacting with miRNAj, M is the number of mRNAs interacting with miRNA_*j*_, *n* is the number of the shared mRNAs by miRNA_*j*_ and miRNA_*j*_

The miRNA-miRNA pairs with significant sharing of mRNAs (e.g. p-value < 0.05) are regarded as miRNA-miRNA synergistic pair. We set the p-value cutoff as 0.05 (adjusted by Benjamini & Hochberg method). As shown in Fig 7, each miRNA tends to synergistically regulate target genes with at least one other miRNA. In terms of its top 50, 100, 150 and 200 target genes, each miRNA synergistically regulates target genes with at least 9, 11, 10 or 11 other miRNAs, respectively. This result indicates that miRNAs may involve in many biological processes by synergistically regulating target genes.

**Fig 7.**
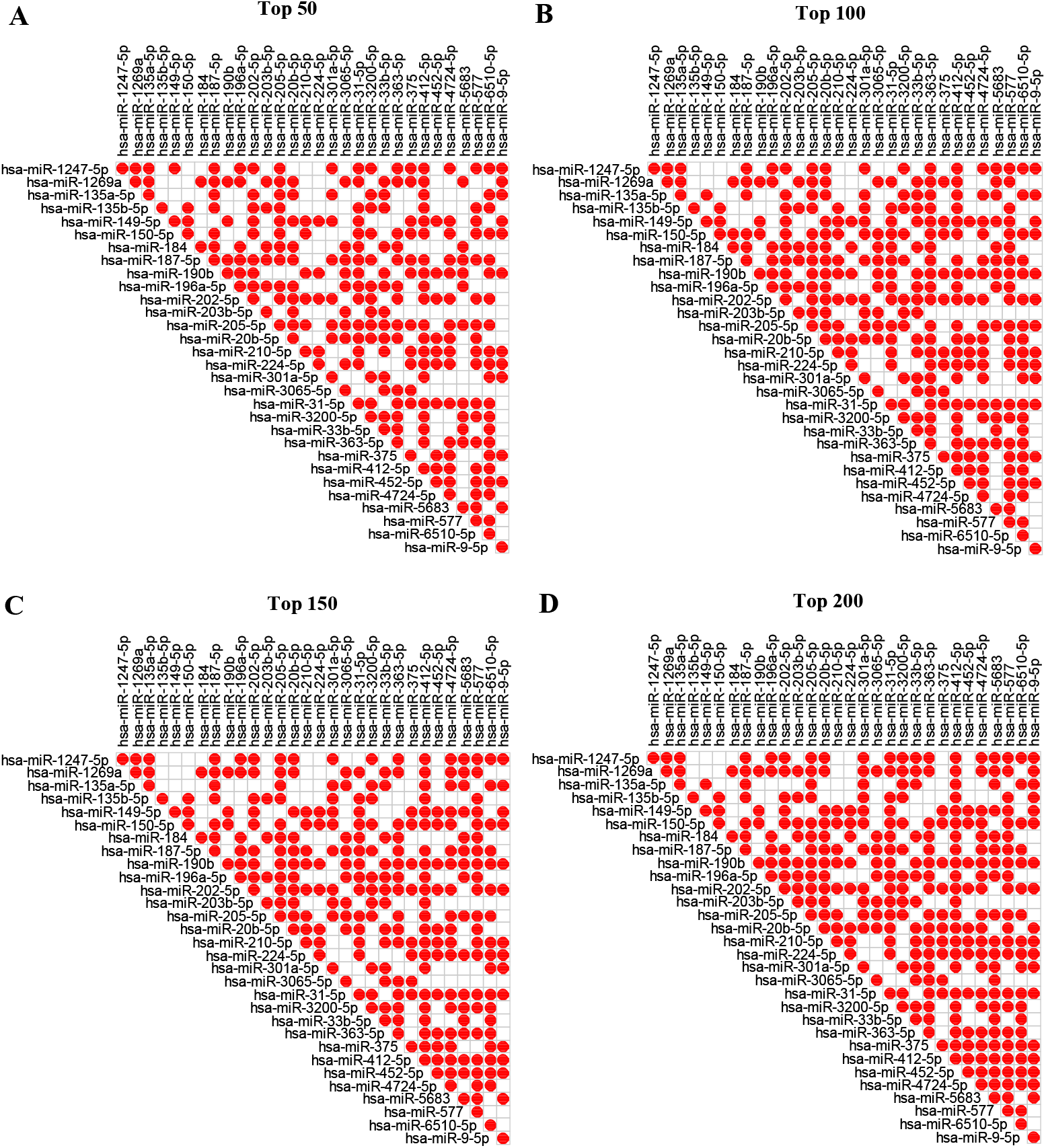
Heatmap of miRNA-miRNA synergistic relationships. Relationships in the top 50, 100, 150 and 200 target genes for each miRNA identified by hiddenICP-Pam50. A red dot indicates that there is a synergistic relationship between two miRNAs.

### Several miRNAs are significantly enriched in functions or diseases related to BRCA

In this section, we conduct GO [24], KEGG [25], Reactome [26] and DisGeNET [27] enrichment analysis of top target genes for each miRNA identified by hiddenICP-Pam50. Since the enrichment analysis results of hundreds of target genes are too general to gain biological insight, we only focus on the enrichment analysis of the top 50 target genes for each miRNA. In Table 1, out of the 30 miRNAs, 12, 10, 13 and 18 miRNAs are significantly associated with at least one GO, KEGG, Reactome and DisGeNET terms, respectively. As shown in Table 2, several miRNAs are significantly enriched in functions or diseases related to BRCA. The results show that the findings using our methods are biologically meaningful in the BRCA dataset. The detailed enrichment analysis results can be seen in S6 File.

**Table 1.** Functional enrichment analysis of the top 50 target genes for each miRNA identified by hiddenICP-Pam50 (at least one term more than 0).

| miRNAs | #GO terms | #KEGG terms | #Reactome terms | #DisGeNET terms |
| --- | --- | --- | --- | --- |
| hsa-miR-187-5p | 3 | 0 | 0 | 0 |
| hsa-miR-1269a | 7 | 0 | 0 | 0 |
| hsa-miR-184 | 0 | 1 | 0 | 0 |
| hsa-miR-205-5p | 23 | 0 | 2 | 147 |
| hsa-miR-196a-5p | 7 | 8 | 0 | 58 |
| hsa-miR-203b-5p | 2 | 1 | 2 | 0 |
| hsa-miR-375 | 0 | 2 | 5 | 23 |
| hsa-miR-5683 | 33 | 0 | 0 | 54 |
| hsa-miR-6510-5p | 0 | 1 | 11 | 0 |
| hsa-miR-20b-5p | 2 | 0 | 0 | 1 |
| hsa-miR-363-5p | 51 | 1 | 4 | 56 |
| hsa-miR-577 | 0 | 0 | 2 | 38 |
| hsa-miR-135b-5p | 27 | 0 | 0 | 26 |
| hsa-miR-150-5p | 17 | 0 | 3 | 32 |
| hsa-miR-412-5p | 0 | 0 | 0 | 25 |
| hsa-miR-1247-5p | 40 | 0 | 15 | 8 |
| hsa-miR-4724-5p | 0 | 0 | 5 | 0 |
| hsa-miR-3065-5p | 0 | 0 | 0 | 4 |
| hsa-miR-31-5p | 1 | 2 | 0 | 1 |
| hsa-miR-135a-5p | 0 | 0 | 2 | 2 |
| hsa-miR-9-5p | 0 | 2 | 5 | 32 |
| hsa-miR-3200-5p | 0 | 1 | 0 | 21 |
| hsa-miR-33b-5p | 0 | 0 | 4 | 0 |
| hsa-miR-202-5p | 0 | 5 | 0 | 39 |
| hsa-miR-301a-5p | 0 | 0 | 3 | 6 |

**Table 2.**
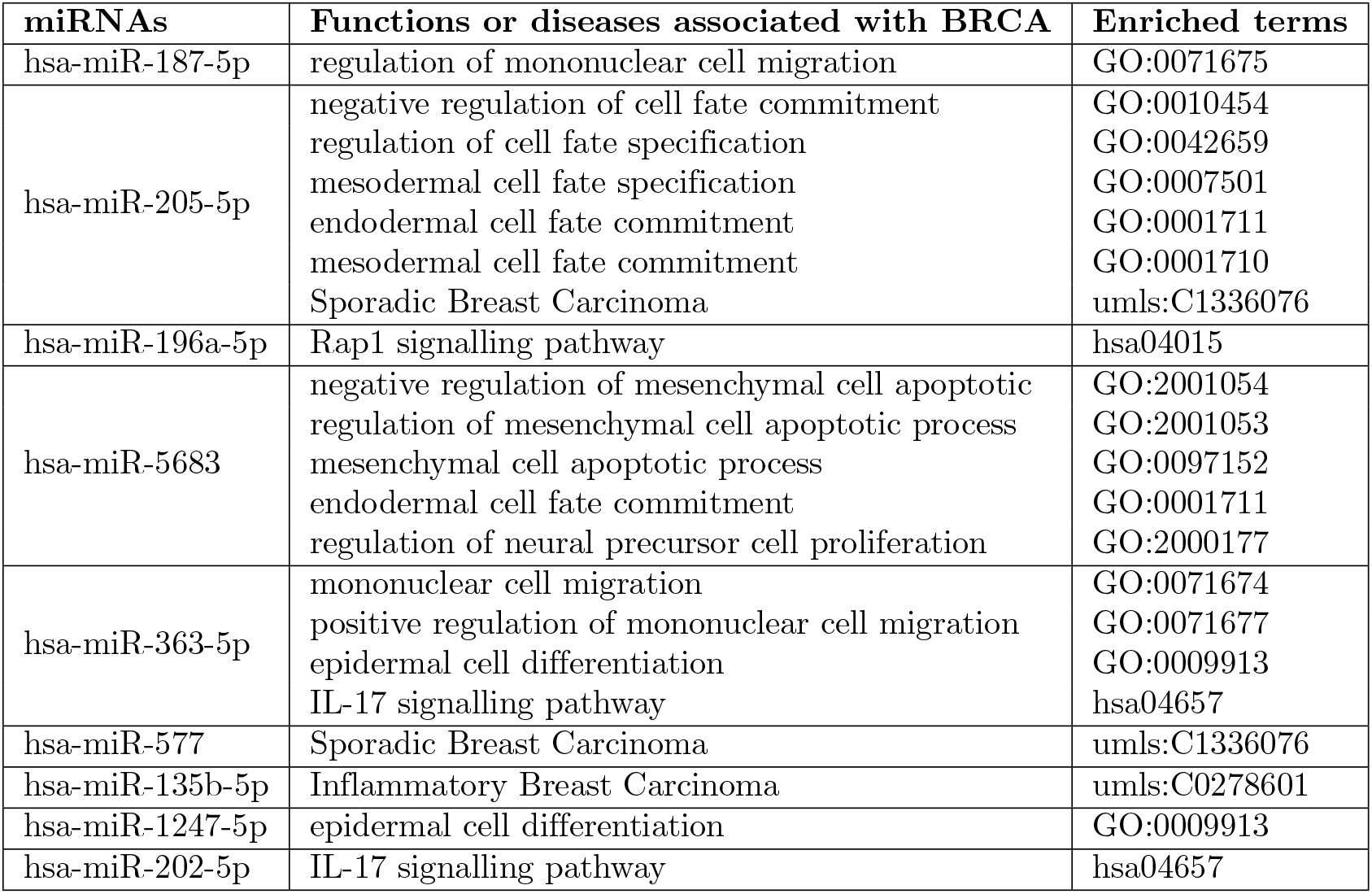
Several miRNAs are significantly enriched in functions or diseases related to BRCA.

## Conclusion

From the fact that miRNAs regulate gene expression by binding the 3’-UTR of mRNAs at the post-transcriptional level [6,35–37], they are important in various biological processes in the human body and identifying their regulation mechanisms plays a salient role in diagnostics and therapeutics for a wide range of diseases. At the present, although numerous studies have developed methods to identify the relationships of miRNAs and mRNAs, most of them detect the correlations between the expression levels of miRNAs and mRNAs while the methods discovering the cause-effect relationship have a high computational complexity. To deal with this problem, we introduce the methods to identify causal effects of miRNAs on mRNAs based on ICP [20].

ICP is a method which is used to infer causality of variables across different environments such as different datasets obtained from different sources/labs for studying the same disease or different types of datasets (observational data and data obtained from intervention experiments), and it is based on the invariance assumption of the causal relationships across different settings. The method has been designed with high dimensional data in mind and has an extension for hidden variables. These features have made the ICP method a great candidate for dealing with biological problems, where the datasets (such as gene expression data) may contain measurements of thousand of variables while some variables are hidden/unobservable.

For our method, first of all, we select top miRNAs and mRNAs with the most different median absolute deviation from BRCA dataset. We then apply Pam50 method to categorise BRCA samples into different environment settings based on different cancer subtypes. After that, we use the invariant causal prediction to find miRNA-mRNA regulatory relationships across subtypes. We validate the results with the miRNA-transfected experimental data and the results show that our method outperforms others. Moreover, to take the advantages of hiddenICP as well as Pearson and Lasso, we combine them into the ensemble method using Borda election to discover miRNA-mRNA regulatory relationships. We validate the results with the experimentally confirmed data and it shows that our algorithm outperforms other methods in finding the interactions and can complement to other methods in finding miRNA-mRNA interactions. Further enrichment analysis indicates that miRNAs involved in the predicted regulatory relationships tend to synergistically regulate target genes, indicating the usefulness of our methods in uncovering miRNA regulation mechanisms.

## Supporting information

**S1 File. BRCA data set.** The expression of matched miRNAs and mRNAs of the breast adenocarcinoma (BRCA) data set is downloaded from The Cancer Genome Atlas (TCGA).

**S2 File. Top 2,000 interactions for all miRNAs.** Top 2,000 predicted miRNA-mRNA interactions for all miRNAs by hiddenICP-Pam50.

**S3 File. Top 50, 100, 150 and 200 interactions for each miRNA.** Top 50, 100, 150 and 200 predicted miRNA-mRNA interactions for each miRNA by hiddenICP-Pam50.

**S4 File. The transfection data.** The transfection data for validating the predicted results of miRNA-mRNA regulation relationships.

**S5 File. The experimentally confirmed databases.** The confirmed miRNA-mRNA interactions retrieved from miRTarbase 6.1, TarBase 7.0, miRWalk 2.0.

**S6 File. The detailed enrichment analysis results.** The detailed enrichment analysis results of functional annotation of miRNAs.

## Acknowledgments

The NHMRC Grant (No: 1123042); the Australian Research Council Discovery Grant (No: DP170101306); the National Natural Science Foundation of China (No: 61702069); the Applied Basic Research Foundation of Science and Technology of Yunnan Province (No: 2017FB099); Presidential Foundation of Hefei Institutes of Physical Science, Chinese Academy of Sciences (No. YZJJ2018QN24); and Australian Government Research Training Program Scholarship.

